# Topological Data Analysis of Spatial Protein Expression in Multiplexed Spatial Proteomics Studies

**DOI:** 10.64898/2026.02.25.707521

**Authors:** Sarah Samorodnitsky, Michael C. Wu

**Affiliations:** Public Health Sciences Division, Fred Hutchinson Cancer Center; SWOG Statistics and Data Management Center, Fred Hutchinson Cancer Center

## Abstract

Multiplexed spatial proteomics platforms generate high-resolution images capturing the spatial expression of proteins in tissue. Images are often fed through a complex pre-processing pipeline to identify individual cells (termed segmentation) and then to predict their phenotypes. It is common to test if the inferred spatial arrangement of cells associates with patient-level outcomes. However, cell segmentation and phenotyping are prone to error and this approach neglects the measured protein levels. Further, new research suggests topological analysis of spatial proteomics may yield more power than alternative approaches. We propose a method, `TOASTER`, that circumvents reliance on segmentation and phenotyping and instead tests the association between continuous spatial protein expression and a patient-level response variable. `TOASTER` uses topological data analysis to first characterize the presence of topological features within univariate and bivariate spatial protein expression. The topological structure is summarized using an adaptation of the Nelson-Aalen cumulative hazard function. We can then associate this summary with an outcome using either a functional data analytic approach, a gridwise testing approach, or using kernel association testing. We show via simulation that our approach improves power and controls type I error, even in the presence of gaps or tears in the image which may arise during tissue handling. We apply our approach to a study in triple-negative breast cancer and demonstrate topological features of protein expression associated with immunotherapy response.

## 1 Introduction

Multiplexed spatial proteomics imaging platforms capture the spatial context of protein expression in tissue, enabling study of the relationship between tissue-level protein architecture and patient-level outcomes [1]. Many popular spatial proteomics platforms, such as multiplexed ion beam imaging (MIBI) [2], imaging mass cytometry (IMC) [3], and multiplexed immunohistochemistry [4], use fluorescence- or metal-tagged antibodies to quantify a targeted set of proteins whose locations are recorded via high-resolution imaging. The locations and phenotypes of individual cells can be identified based on these images, providing a snapshot of the cellular activity within tissues. The inferred spatial organization of cells can then be associated with patient-level outcomes. This has led to clinically meaningful discoveries such as how spatial patterns among cells are altered by immunotherapy [5] and whether spatial interactions between tumor and immune cells associates with survival [6].

Raw spatial proteomics data often comes in the form of multichannel TIFF images where each channel contains the intensity of the unique antibody used to detect a particular protein [7]. Standard analysis pipelines require raw images to undergo complex pre-processing which includes artifact removal (e.g., autofluorescence), segmentation (prediction of the boundaries of individual cells and nuclei), and phenotyping (prediction of the cell type labels) [8]. From there, the images are often analyzed as realizations of a marked spatial point process [9] wherein each cell is treated as a point with an associated “mark” denoting its cell type, e.g. a CD8+ T cell or B cell [7]. Many methods have been developed to test the association between these marked spatial point patterns and patient-level outcomes [10, 11, 12, 13, 14, 15].

A significant challenge in using the spatial point process model is that it relies on the predicted boundaries and phenotypes assigned to cells during pre-processing. Overlapping cells, varying or uneven cell morphologies, and sliced cells can induce error in the segmentation and phenotyping results [8]. In addition, protein expression beyond predicted cell boundaries may be discarded, losing meaningful information about the tissue [16]. Finally, by focusing only on cell types, the expression and intensities of the individual proteins is ignored. These challenges in the analysis of multiplexed spatial proteomics data have not been addressed in the existing statistical methods literature.

An alternative approach is to avoid cell segmentation and phenotyping and instead analyze the continuous, spatial protein expression captured by the multiplexed imaging platforms. Cell segmentation and phenotyping rely on the correlation between protein expression and cell type. For example, if the CD3 protein is highly expressed within the predicted boundaries of a cell, phenotyping algorithms may predict that that cell is a T cell. However, we may similarly interpret that a tissue region exhibiting elevated CD3 expression contains T cells (and, conversely, a region with a lack of CD3 expression is absent of T cells). Thus, analyzing continuous spatial protein expression may yield similar (or unique) insights into the composition of the tissue while circumventing reliance on imperfect cell segmentation and phenotyping output. However, relatively little has been done in this area, and few statistical methods are available to associate the continuous spatial protein expression with patient-level outcomes.

Given the limitations of current spatial point process-based approaches and the shortage of strategies for analyzing spatial protein expression, we propose `TOASTER` (**T**est **O**f **A**ssociation between **S**patial protein expression and clinical **T**raits-of-int**ER**est) which is a global test of association between continuous, spatial protein expression and patient-level outcomes. `TOASTER` uses topological data analysis (TDA) [17] to summarize the topological structures (namely, connected components and loops) present in the spatial protein expression and then tests whether these topological structures are related to the clinical outcome, which may be a continuous, binary, or censored survival endpoint. Operationally, we borrow a recent approach developed for topological analysis of random fields [18] that characterizes the topological structure for each sample as a cumulative hazard function where events are births of new topological structures during filtration. We refer to these summaries as “topological event history,” and adopt this strategy to assess the expression of a single protein (Figure 1) and then further develop an extension to summarize topological structures shared between pairs of proteins. We can then test if the topological event history is associated with a clinical outcome and propose three different approaches based on treating the cumulative hazards as functions, considering a grid of function values, and using a kernel machine testing approach [19].

**Figure 1.**
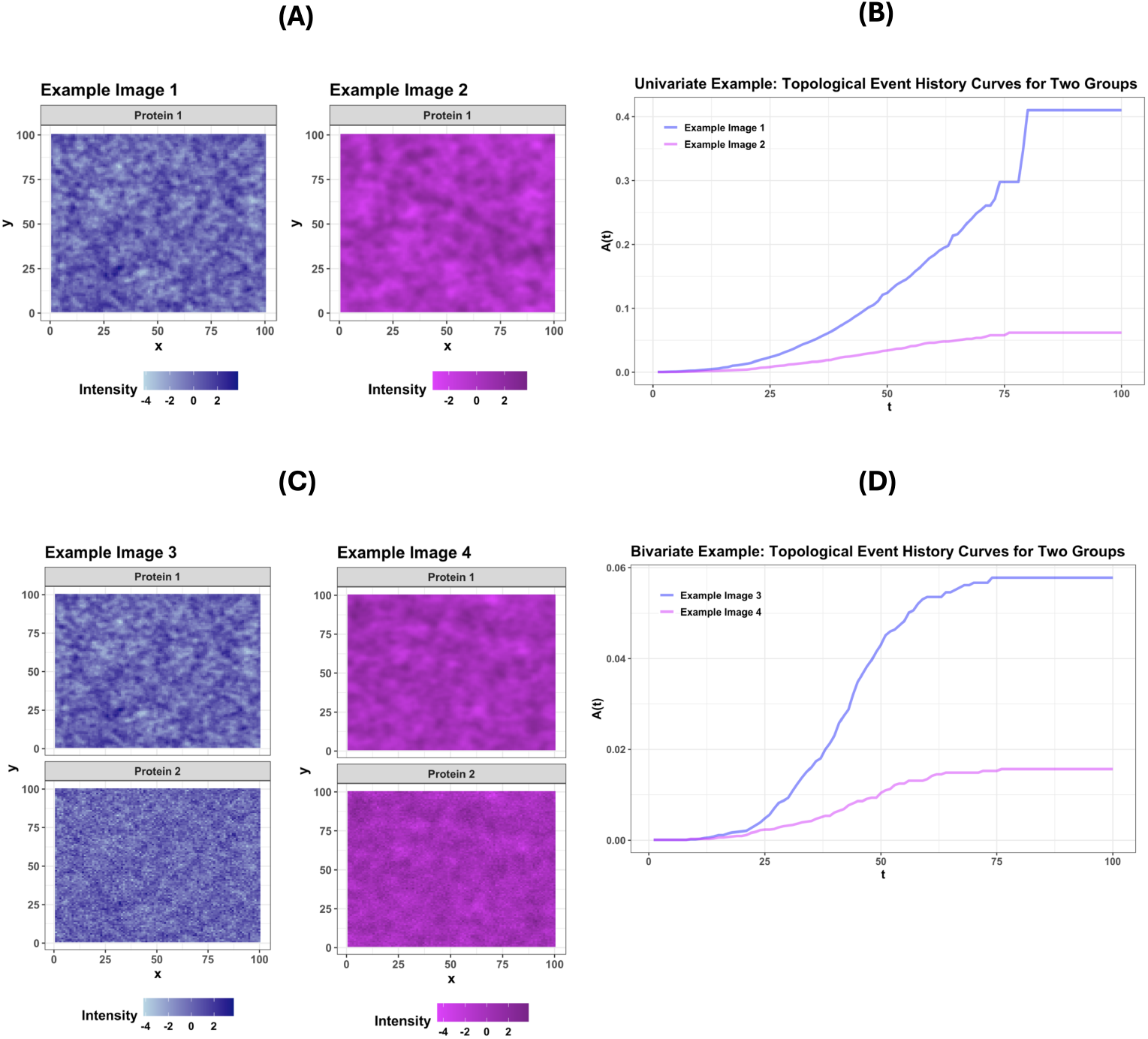
Topological event history of univariate and bivariate simulated spatial protein expression. (A) Two simulated protein expression patterns with differing correlation structures. (B) Topological event history functions for patterns in (A). (C) Two simulated bivariate spatial protein expression patterns. (D) Topological event history functions for patterns in (B).

The primary advantage of `TOASTER` is that it treats protein expression as continuous. This avoids the reliance on cell segmentation and phenotyping and also offers improved power in situations where quantitative expression levels are related to outcomes. Practically, `TOASTER` allows for holes or tears in tissue samples that may occur during sample processing, a frequent challenge in the analysis of spatial proteomics imaging [7].

The rest of this paper is organized as follows. In Section 2, we introduce `TOASTER` in the context of a single protein marker. We then present the adaptation necessary to accommodate two protein markers before describing three strategies for testing the association between topological structure in the spatial protein expression and patient-level phenotypes. In Section 3, we benchmark the performance of `TOASTER` against existing methods in a simulation study. In Section 4, we discuss an application of `TOASTER` to a study of combination chemotherapy and immunotherapy in triple negative breast cancer. Finally, we discuss some future directions in Section 5.

## 2 Methods

### 2.1 Notation

Suppose we have an image (e.g. a tumor biopsy) from each of *n* subjects. The dimension of each image is denoted *d*_*i*1_ *× d*_*i*2_ for *i* = 1, …, *n* and may vary across samples. Multiplexed spatial proteomics data is often stored in multichannel TIFF images, where each channel contains the detected expression of each protein/antibody marker at that location [7]. Note that we use the terms “protein” and “marker” interchangeably throughout this text. Assume that we have probed the expression of *M* proteins. Within each multichannel TIFF image, a vector 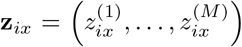 denotes the expression of the *M* proteins at location *x* where *x* = 1, …, *d*_*i*1_*d*_*i*2_. Higher values mean greater pixel intensity and therefore higher protein expression. We let 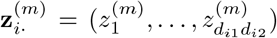 denote the expression for the *m*^*th*^ marker for subject *i* across the image. We assume that for each subject the intensity of each protein has been centered and standardized. The goal is to describe the topological structure of either a single marker (each 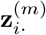) or two markers jointly (i.e. an interaction between the *m*-th and *m*^*′*^-th markers, 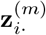 and 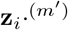) and then test if this structure across images associates with patient-level outcomes. The topological structures we wish to capture are degree-0 homologies, which are connected components, and degree-1 homologies, which are loops. A comprehensive overview of topological data analysis is provided in [20].

In what follows, we first introduce the methodology for describing the topological structure of a random field (representing the spatial expression of a single protein) in Section 2.2 and its adaptation and contextualization for spatial proteomics data. We then discuss extending this approach to describe the shared topological structure of two markers in Section 2.3, which may represent two proteins that are often expressed on the surface of the same cell types or the colocalization of two cell types. We then describe three approaches (a functional approach, a gridwise approach, and a kernel approach) for testing whether the topological structure of our images associate with sample-level outcomes in Sections 2.4.1, 2.4.2, and 2.5, respectively. We temporarily drop the subscript *i* for ease of explanation.

### 2.2 Univariate TOASTER

Given a single marker *m*, we use the approach of [18] in characterizing the topological structure of spatial protein expression within a tissue sample and summarizing this structure using topological event history. We first review this approach for tracking the births of connected components in a random field, with the understanding that tracking loops can proceed similarly and is described further below.

To detect connected components, we apply a filtration to each image. The filtration iteratively thresholds the intensity of marker *m* in the image at level *t*. We start by setting *t* such that no pixels have intensity below the threshold. Then, we increment the threshold such that *t* exceeds the intensity of some pixel. This pixel is said to appear or to be “born.” We continue increasing *t* and track the appearance (i.e, birth) of pixels who are not adjacent to any already-born neighbors. These pixels represent connected components in the field. During filtration, the birth of connected components reveals “new” regions of the field that had not yet appeared. We do this until all pixels have been born. A visualization of this process for a single protein is provided in Figure 2.

**Figure 2.**
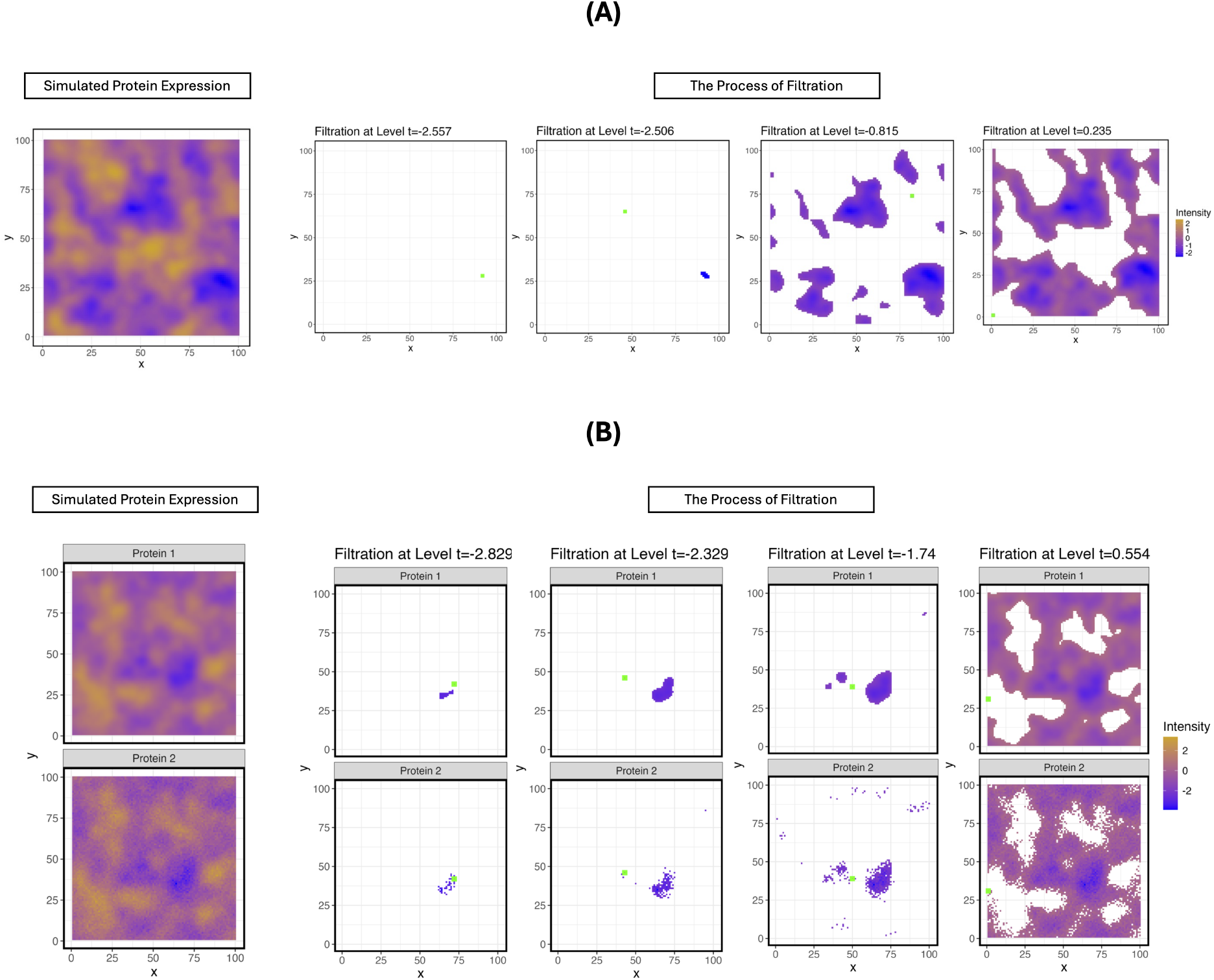
(A) An example of applying the filtration to a single protein. The full simulated image of protein expression is shown on the left hand side. On the right hand side, four steps during the filtration process are shown. The pixel born at each step is highlighted in green. (B) An example of applying the filtration to two proteins. The full simulated image of bivariate protein expression is shown on the left hand side. On the right hand side, four steps during the filtration process are shown. The pixel born at each step is highlighted in green.

We then treat the birth of each new connected component as an “event” that occurs at threshold *t*. Then, the sequence of births can be summarized using survival analysis techniques. Specifically, we define an event having occurred at location *x* at threshold *t* if:

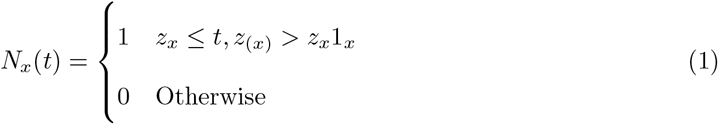

where *z*_*x*_ represents the value of the random field (the intensity of the protein) at location *x, z*_(*x*)_ is the vector of intensity values of the neighboring pixels of *x*, and 1_*x*_ is a vector of 1s the same length as *z*_(*x*)_ (i.e., the number of pixels immediately adjacent to location *x*). In other words, an event occurs at location *x* if *x* is a local minimum or has value, *z*_*x*_, smaller than its neighbors, *z*_(*x*)_.

At threshold *t*, a location *x* is considered to be at-risk if:

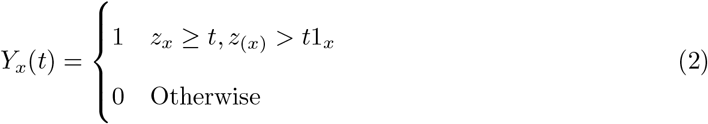

When *Y*_*x*_(*t*) = 1, location *x* remains at risk. A pixel leaves the risk set if it is born as a local minimum by level *t* or if one of its neighbors is born as a local minimum by level *t*.

We let *N* (*t*) = ∑_*x*_ *N*_*x*_(*t*), which counts the total number of events that occurred at *t*, and *Y* (*t*) = ∑_*x*_ *Y*_*x*_(*t*), which counts the total number of pixels still at risk at *t*. Then we summarize the births of connected components throughout filtration using an adaptation of the Nelson-Aalen cumulative hazard:

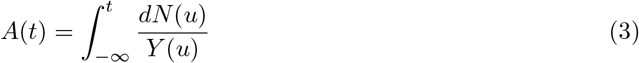

which we estimate via:

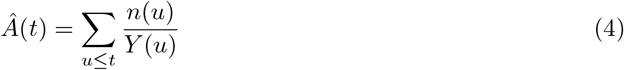

where *n*(*u*) is the instantaneous number of events that occurred at *u*. Pseudocode to calculate *Â*(*t*) is given in Algorithm 1. *Â*(*t*) is referred to as the “topological event history” of the given protein and summarizes the topological structure of its spatial context. We can then assess the topological event history functions across images for associations with clinical outcomes.

#### Algorithm 1

Tracking births of local minima in spatial expression of a single protein within a sample (Reproduced from the software provided by [18])

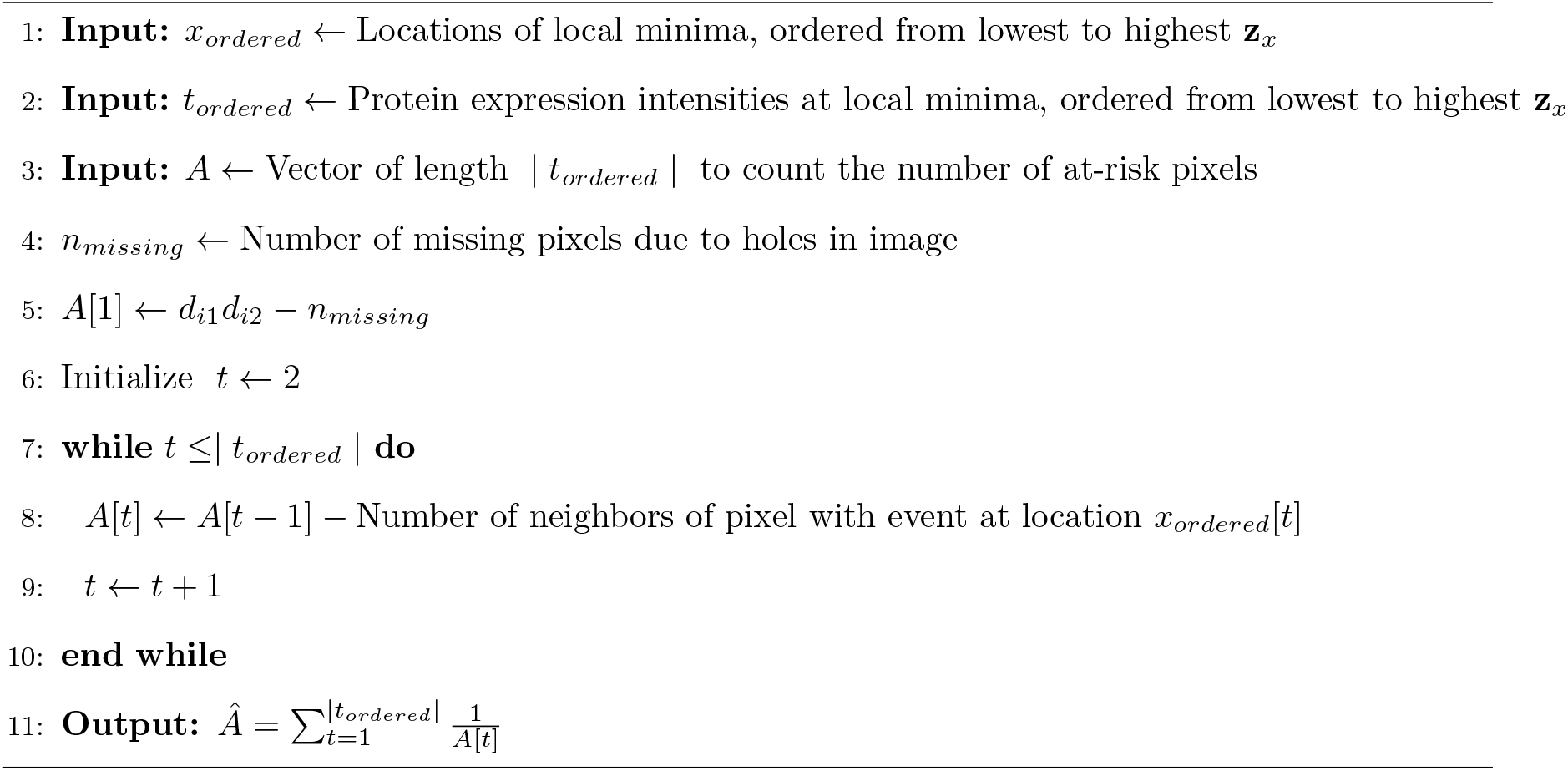

For our analyses, neighbors of a location are determined “rook-wise,” i.e. the four pixels above, below, to the left, and to the right. “Queen-wise” neighbors which further include the diagonal pixels can also be used. Practically, if a pixel lies on the edge of the image, or adjacent to a hole or tear in the image, its set of neighbors is reduced by the number of missing pixels. In doing this, `TOASTER` naturally accommodates gaps in the image, which may arise during handling of the tissue specimens.

Although we have focused on connected components, this approach can be naturally extended to summarize loops and rings in the protein expression. Specifically, we note that the Alexander duality [21] can be exploited to track deaths of loops (rather than births). Thus, to use deaths of loops as events, we simply flip the sign of our random field and track the local minima of the sign-reversed field. In this situation, we are tracking loops that are “filled” or die during the process of filtration on the original scale and the birth of loops on the flipped scale. From there, the same definitions of an event given in Equation 1 and at-risk given in Equation 2 apply and we can use the Nelson-Aalen cumulative hazard function given in Equation 3 as our summary statistic.

### 2.2 Bivariate TOASTER

Beyond considering individual proteins, investigators are often interested in simultaneously analyzing pairs of proteins to study their interactions and associate the interactions with outcomes. We then extend the approach of [18] to the bivariate scenario. This allows us to capture the topological structure in the spatial distribution of a cell type defined by two markers (e.g., CD8 T cells are detected based on CD3 and CD8 expression) or to capture the colocalization between two cell types each defined by a single marker (e.g., T cells are identified based on CD3 and B cells based on CD20). We again focus on connected components with the understanding that the approach can be exploited to study rings, as well.

In considering a pair of proteins, we view their joint spatial expression as a bivariate random field with two “layers,” where each layer corresponds to a single protein. Because the protein expression is measured on the same sample, the pixels across the fields align such that location *x* now has two marker values: 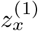 and 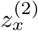 corresponding to markers 1 and 2. We again assume that each layer has been standardized.

In the bivariate case, each location *x* may exhibit one of the following cases:

1. pixel *x* is a local minimum in both layers 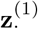 and 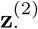
2. pixel *x* is a local minimum in only one layer, 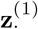 or 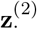
3. pixel *x* is not a local minimum in either layer

We treat Case (1) as an event, but not Cases (2) or (3) because we are interested in situations where the structures correspond. Letting *t*^(1)^ and *t*^(2)^ denote the threshold parameters for the filtration for markers 1 and 2, we define an event at location *x* as:

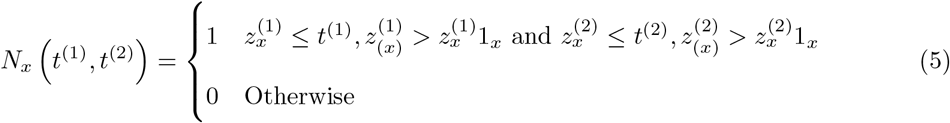

where 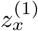 is the value of marker 1 at location *x*, 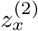 is the value of marker 2 at location *x*, 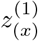 is the value of marker 1 of the neighbors of location *x*, and 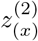 is the value of marker 2 of the neighbors of location *x*.

To determine if location *x* as at-risk, we define an indicator, *T*_*x*_ where *T*_*x*_ = 1 if *x* is a local minimum in both layers (Case (1)) and *T*_*x*_ = 0 if *x* is either a local minimum in only one layer or in neither layer (Cases (2) and (3)). We then define a pixel as at-risk if:

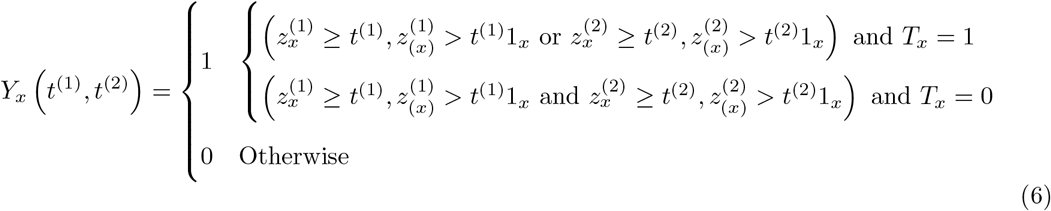

Denoting 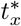 as the threshold at which *x* leaves the risk set, if a location is born as a connected component in both layers (i.e., Case (1)), then 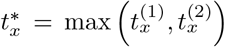 where 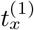 and 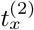 are the respective timings of the births in marker layers 1 and 2. Under Case (2), 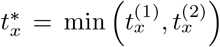 where 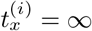 for marker layer *i* where *x* is not a local minimum. Under Case (3), 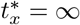. Thus, locations that are not local minima in both layers leave the risk set once they have their terminal event or once their neighbor has their terminal event, whichever comes first. Only locations that are local minima in both layers count as events. This process is visualized in Figure 2 and the algorithm to construct *Â*(*t*) for an image is given in Algorithm 2.

Similar to the univariate case, we then construct a summary tracking the births of shared connected components using the Nelson-Aalen cumulative hazard. We track how many locations are at risk at *t* and count up the events at each terminal timepoint *t*^∗^ when shared local minimum leave the risk set:

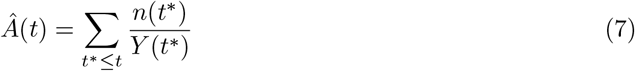

As described in Section 2.2, this process may be analogously applied to track the death of loops by flipping the sign of both fields and tracking the birth of local minima in the sign-flipped field.

#### Algorithm 2

Tracking births of shared local minima in spatial expression of two proteins for a given sample

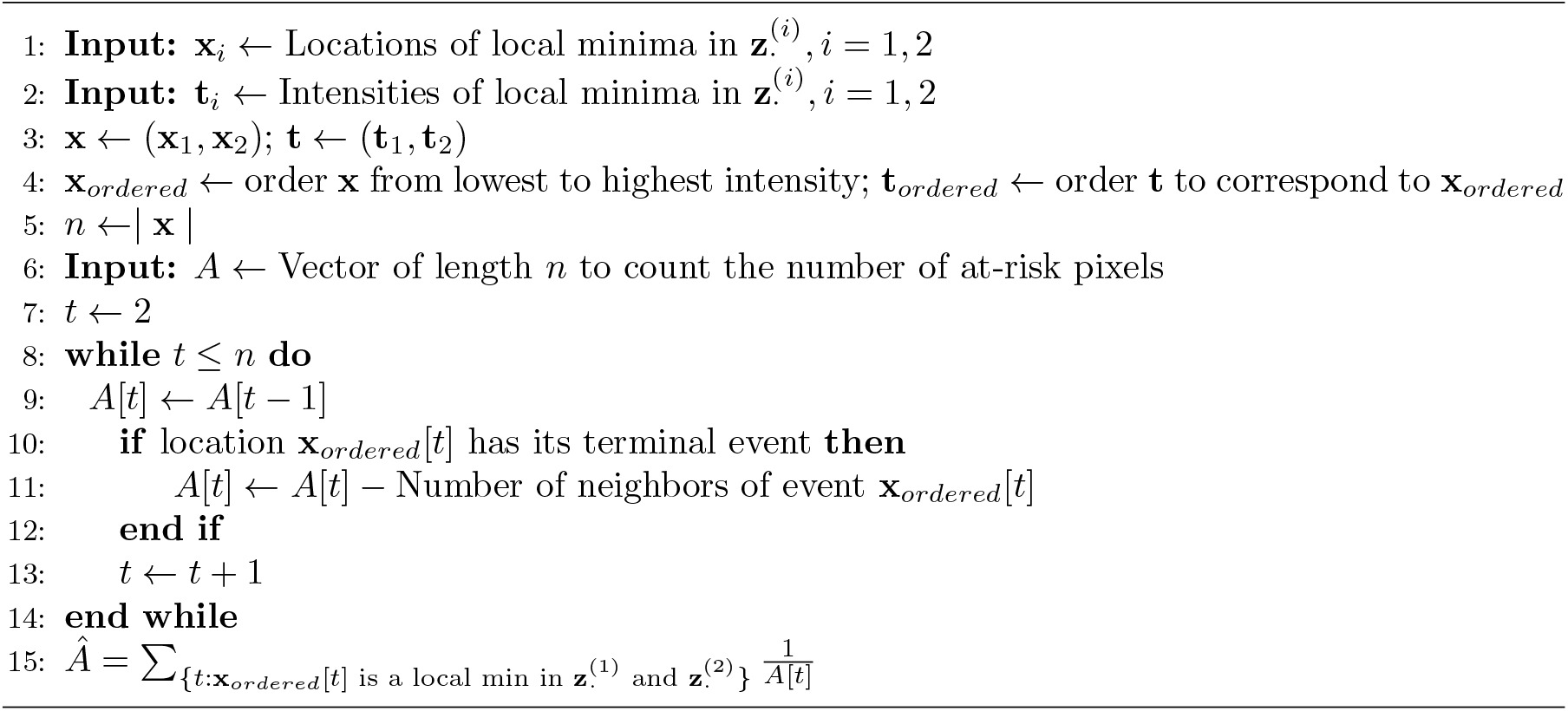

### 2.4 Associating Topological Event History and Response Variables

The objective is to not only to estimate the topological event history of each sample but also to test its association with outcomes. To achieve this, we consider three different approaches. To facilitate each approach, we first align the topological event history curves. For each curve, 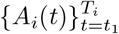, the number of events during filtration, *T*_*i*_, may vary from sample to sample. We thus discretize these curves to the same set of grid points, {*t*_1_, …, *t*_*N*_} to yield 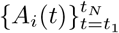. There are many strategies to choose {*t*_1_, …, *t*_*N*_}, but in our simulation study and data application we use 100 quantiles between 0.1 and 0.9.

#### 2.4.1. Association Testing via Functional Analysis

The first testing approach is based on functional principal components analysis (FPCA) [22] which we use to decompose the variation among the topological event history curves into a set of latent functional principal components. Following previous work [14, 23], we treat the FPCA scores for each functional principal component as fixed covariates in an outcome model suitable for the dependent variable. Here, we present our approaches for a right-censored survival outcome and a binary outcome.

Using the Karhunen-Loève expansion, we can represent each curve as:

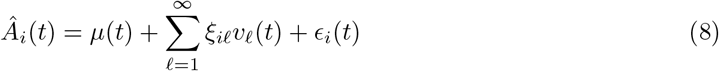

where *µ*(*t*) denotes the overall mean function, *v*_*ℓ*_(*t*) denotes the *ℓ*th eigenfunction, *ξ*_*iℓ*_ denotes the functional principal component scores with 𝔼 (*ξ*_*ji*_) = 0 and Var(*ξ*_*ji*_) = *λ*_*j*_ where *λ*_*j*_ is the eigenvalue corresponding to the *j*th eigenfunction, and *ϵ*_*i*_(*t*)) represents random error with 𝔼 (*ϵ*_*i*_(*t*)) = 0 and Var(*ϵ*_*i*_(*t*)) = *σ*^2^. For a survival outcome, which is used in our simulation study in Section 3, we assume to have event times *T*_*i*_ and censoring times *C*_*i*_ for each patient *i* where *s*_*i*_ = min(*T*_*i*_, *C*_*i*_). We let Δ_*i*_ = 1(*T*_*i*_ ≤ *C*_*i*_) denote whether patient *i* experienced the event. We then use a Cox proportional hazards model to model the relationship between *ξ*_*iℓ*_, *i* = 1, …, *n* and *ℓ* = 1, …, *L*, and the hazard:

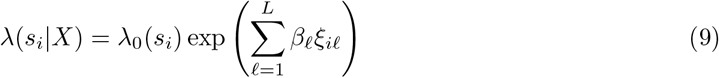

We can further adjust for clinical or demographic covariates, such as age or sex, within the model. We then use a likelihood ratio test to evaluate the global significance of the *ξ*_*iℓ*_s. We include the scores from all functional principal components which explain more than 99% of variation, but different thresholds may be chosen to suit.

For a binary outcome, which we considered in our triple negative breast cancer application in Section 4, we use a logistic regression model to test the relationship between the topological event history curves and the log-odds ratio. Assuming we have a binary outcome *y*_*i*_ for each sample, we model:

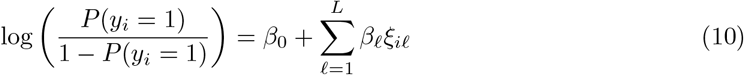

We then use a deviance test where the test statistic is calculated as the difference between the deviance of the fitted model and null model. This statistic follows a *χ*^2^ distribution with *L* degrees of freedom, which is used to calculate the p-value.

#### 2.4.2. Association Testing via Gridwise Testing

We next consider a gridwise approach to testing the association between 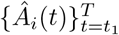 and an outcome. For each *t*_*j*_ ∈ {*t*_1_, …, *t*_*N*_}, we subset the topological event history curves to their values at *t*_*j*_, *Â*_*i*_(*t*_*j*_). We test for an association with the outcome given level *t*_*j*_ using a Wald test resulting from treating the *Â*_*i*_(*t*_*j*_) as a covariate in a Cox proportional hazards model (for a survival outcome) or using a Wilcoxon rank-sum test (for a binary outcome). Applying this to each grid point, we obtain a vector of *N* p-values, {*p*_1_, …, *p*_*N*_} which we combine using the Cauchy combination test [24]. Given *N* p-values, the Cauchy combination test statistic is:

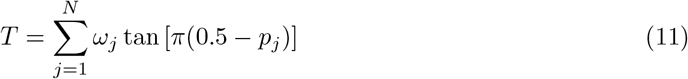

where we fix the weights *ω*_*j*_ = 1*/N* for all *j*. Under the null, *T* follows a mixture of Cauchy distributed random variables. The advantage to using the Cauchy combination test is that it is appropriate for combining dependent p-values and is computationally efficient if a large number of grid points are considered.

### 2.5 Association Testing via Kernel Testing

Finally, we consider a kernel association testing approach. To implement this, we construct an *n × n* pairwise distance matrix of the Euclidean distances between each discretized topological event history curve, i.e.:

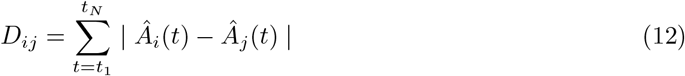

yielding a distance matrix, 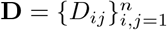. We then convert this to a kernel matrix using a Gower centered kernel:

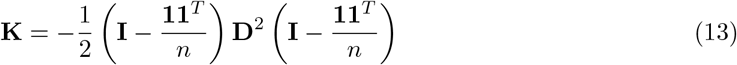

Finally, we use a kernel association testing to examine an association with a response variable. The general form of the kernel association test statistic is:

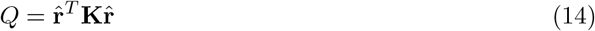

where **K** is the *n × n* kernel matrix and 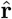 is a vector of residuals from a regression model under the null. Given a survival outcome and a using a Cox model, we use Martingale residuals in our test statistic [25]. For a binary outcome, we use 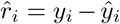 where *ŷ*_*i*_ is the fitted value from the logistic regression model. Note that the residuals are from the models fit under the null, assuming no association between the similarity in topological event history curves and the dependent variable. This allows both for the adjustment of other covariates and also assures that the validity of the test is unaffected even if a poor kernel is chosen.

## 3 Simulation Study

To evaluate the power and type I error rate of `TOASTER`, we simulated spatial protein expression, along with sample-level survival outcomes, and studied the performance of the proposed method in detecting associations between the two. We next describe our simulation study for univariate protein expression (Section 3.1), bivariate protein expression (Section 3.2), and bivariate protein expression with induced holes (Section 3.3).

### 3.1 Univariate Simulation

We generated univariate spatial protein expression within 100 *×* 100 pixel images using Gaussian random field models with marginal Gaussian(0, 1) distributions. Following previous work [18], we used a Matérn correlation function to induce correlation between adjacent pixels within each image:

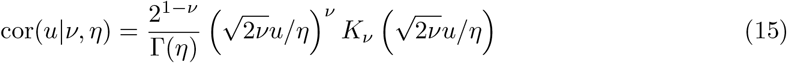

where *u* represents the distance between two pixels and *K*_*ν*_ (·) is a modified Bessel function of the third kind. To generate the data, we split the full sample set of *n* = 100 samples into two groups of size 50 and simulated the images under two different values of parameters (*ν, η*). For the first 50 samples we simulated spatial protein expression using (*ν*_1_ = 5, *η*_1_ = 1). For the latter 50 samples, we simulated spatial protein expression using parameters (*ν*_2_ = 1, *η*_2_ = 1).

For the dependent variable, we simulated time-to-event outcomes from an exponential distribution with parameters *λ*_1_ and *λ*_2_ for the first 50 and second 50 samples, respectively. We varied the hazard ratio *λ*_1_*/λ*_2_ by fixing *λ*_2_ = 1 and varying *λ*_1_ ∈ {2, 1}. We randomly censored 10% of observations in each group.

We used `TOASTER` to track the births of connected components (“`TOASTER` (Degree 0)”) and the deaths of loops (“`TOASTER` (Degree 1)”). Few existing methods are available that are designed for analyzing the spatial protein expression directly, so we compared `TOASTER` with an existing method, DenVar [12], which we adapted for this context. DenVar uses kernel density estimation (KDE) to estimate the marginal distribution of a protein in each image. DenVar then uses the Jensen-Shannon distance (JSD) to compare KDEs between images to construct a distance matrix which is in turn used in a kernel testing framework to test the association between protein expression and a response variable. To apply DenVar, we standardized the marker intensity to be between 0 and 1.

We assessed power across 1000 simulation replications and type I error across 5000 simulation replications. Table 1 shows the power of `TOASTER` vs. DenVar. Across all three outcome modeling types and both connected components or loops, we found `TOASTER` offered higher power than DenVar in detecting an association between the spatial protein expression and survival. We also observed that `TOASTER` controlled type I error at the nominal rate, 0.05, while DenVar was conservative under these conditions. The conservativeness of DenVar may be a result of its adaptation to a context beyond its original design.

**Table 1.** Power and type I error results for associating the spatial expression of a single protein with a survival outcome. We evaluated the performance of `TOASTER` based on tracking connected components (Degree 0) and loops (Degree 1) and using three different outcome modeling approaches (functional modeling, gridwise testing, and kernel association testing).

| Model | Power |  |  | Type I Error |  |  |
| --- | --- | --- | --- | --- | --- | --- |
|  | Functional | Gridwise | Kernel | Functional | Gridwise | Kernel |
| TOASTER (Degree 0) | 0.880 | 0.884 | 0.882 | 0.051 | 0.052 | 0.053 |
| TOASTER (Degree 1) | 0.876 | 0.882 | 0.882 | 0.052 | 0.052 | 0.054 |
| DenVar | 0.490 |  |  | 0.016 |  |  |

### 3.2 Bivariate Simulation

For the bivariate simulation, we again split the full sample set of *n* = 100 samples into two groups of 50. We used the same Gaussian random field model used in the univariate simulation. We generated the first protein with correlation parameters (*ν*_1_ = 5, *η*_1_ = 1) for the first 50 samples and (*ν*_2_ = 1, *η*_2_ = 1) for the second 50 samples. We then simulated the second protein based on the first using the following linear model:

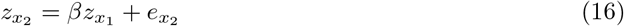

where we varied the value of *β* ∈ {3, 5} and simulated 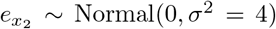. Under *β* = 3 or *β* = 5, the two proteins can be thought of as “colocalizing.” We also considered simulating the two proteins independently (but under the same random field model), in which we simply repeated the same process described in Section 3.1 for both proteins independently. We standardized the intensity of the expression of each protein to be mean 0 and variance 1.

We compared `TOASTER` to an existing method, DIMPLE [11]. DIMPLE is similar to DenVar but estimates the KDE of two proteins and computes the JSD between the KDEs of the two markers within each image. DIMPLE then tests for an association between the JSDs for each image and an outcome using a Cox proportional hazards model. We again simulated a survival outcome as described above and assessed power across 1000 simulation replications and type I error across 5000 simulation replications.

The results are given in Table 2. Across all conditions, we found that `TOASTER` offered slightly higher power than DIMPLE, though both offered high power and controlled type I error. We observed that `TOASTER` performed particularly well when the two markers were simulated independently but from the same parametric random field model. This may be because `TOASTER` captures the differences in the parametric random field used to generate the two groups of images, as previously described [18]. Under this condition, the differences in the parametric random field models used (and the topological structure inherent to these models), rather than the shared topological structure between the two proteins, is associated with the response variable.

**Table 2.** Power and type I error results for associating the spatial expression of a two proteins with a survival outcome.

| Model | $\beta$ | Marker Relationship | Power | | | Type I Error | | |
| --- | --- | --- | --- | --- | --- | --- | --- | --- |
|  |  |  | Functional | Gridwise | Kernel | Functional | Gridwise | Kernel |
| TOASTER (Degree 0) | 5 | colocalize | 0.906 | 0.906 | 0.908 | 0.051 | 0.051 | 0.055 |
| TOASTER (Degree 1) | 5 | colocalize | 0.910 | 0.910 | 0.916 | 0.052 | 0.051 | 0.054 |
| DIMPLE | 5 | colocalize | 0.886 |  |  | 0.052 |  |  |
| TOASTER (Degree 0) | 3 | colocalize | 0.888 | 0.892 | 0.900 | 0.056 | 0.056 | 0.058 |
| TOASTER (Degree 1) | 3 | colocalize | 0.898 | 0.900 | 0.902 | 0.055 | 0.056 | 0.058 |
| DIMPLE | 3 | colocalize | 0.888 |  |  | 0.053 |  |  |
| TOASTER (Degree 0) | 0 | independent | 0.880 | 0.874 | 0.884 | 0.053 | 0.053 | 0.054 |
| TOASTER (Degree 1) | 0 | independent | 0.900 | 0.902 | 0.902 | 0.053 | 0.054 | 0.054 |
| DIMPLE | 0 | independent | 0.712 |  |  | 0.053 |  |  |

### 3.3 Holes

Finally, we present results under the bivariate case with induced random holes in each image. To generate this data, we simulated bivarate spatial protein expression as described in Section 3.2 and then generated two centers uniformly across the field. We then removed the marker expression from any pixel within 5 units from the hole center. The values within the hole were treated as missing. Survival outcomes for each sample were generated as described above. For this simulation scenario, we again compared against DIMPLE which is suitable for use even in the presence of holes in the images [11].

The results are given in Table 3. As in Section 3.2, `TOASTER` offered slightly higher power than DIMPLE when the two proteins were colocalized but significantly improved power when the two proteins were simulated independently. Both approaches controlled type I error at the nominal rate.

**Table 3.**
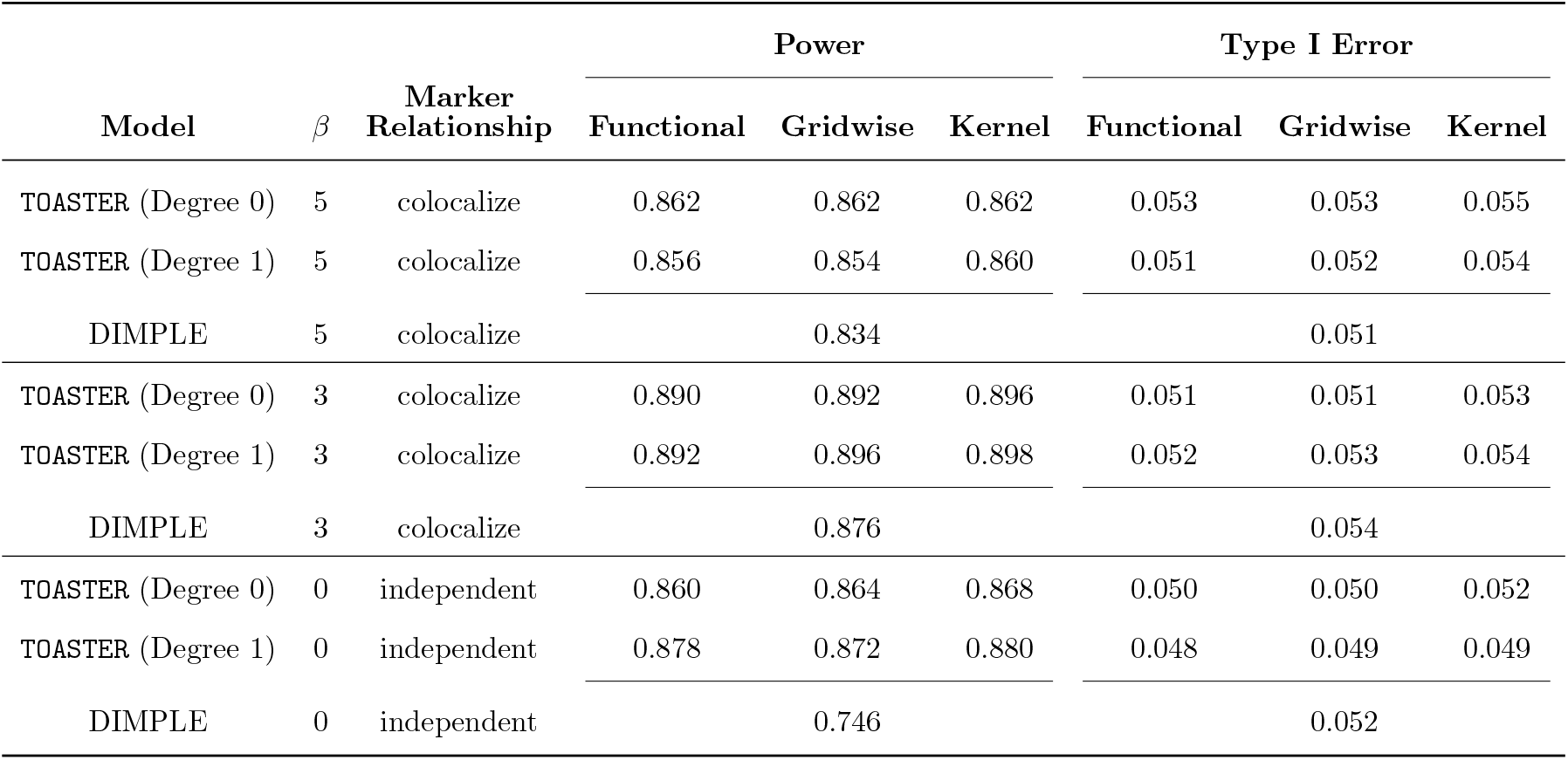
Power and type I error results for associating the spatial expression of a two proteins with a survival outcome in the presence of holes inducing missing data in each image.

## 4 Application to Triple Negative Breast Cancer

We used `TOASTER` to analyze imaging mass cytometry (IMC) data from the NeoTRIP study, which investigated the impact of immune checkpoint blockade (ICB) in patients with triple negative breast cancer [5]. The study compared outcomes in those who received combination ICB plus chemotherapy vs. patients who received chemotherapy alone. IMC was used to capture the spatial context of proteins in the tumor microenvironment before and after treatment with the goal of correlating spatial protein expression with patient-level outcomes. For our analysis, we focus on patients in the combination treatment arm and compared the spatial expression of proteins in those who responded vs. those who did not at the post-treatment time point. Treatment response was encoded as a binary variable: either a patient experienced pathologic complete response (pCR) or a patient experienced recurrent disease (RD).

Our goal was to assess whether the topological structure of the CD3, CD4, CD8, and CD20 proteins associated with treatment response. We chose these proteins for their immunologic importance in cancer: CD3 is expressed by T cells, which are a hallmark of the adaptive immune system [26]; CD3 and CD4 are jointly expressed by CD4+ T cells, a subtype of T cell known to be involved in tumor control [27]; CD3 and CD8 are jointly expressed by CD8+ T cells, a T cell subtype stimulated by ICB to produce an anti-tumor immune response [28]; and CD20 is expressed by B cells, whose increased presence is associated with response to immunotherapy [29]. For our analysis, we tracked connected components within CD3 expression and shared connected components within joint CD3 and CD4 expression, joint CD3 and CD8 expression, and joint CD3 and CD20 expression.

To start, we first processed the raw multichannel TIFF images. To facilitate efficient memory usage, we smoothed the images to have a maximum size of 10, 000 pixels while maintaining each image’s aspect ratio. We also standardized the expression of each protein to have mean 0 and standard deviation 1. Example images for two patients, one who exhibited pCR and one who exhibited RD, are shown in Figure 3.

**Figure 3.**
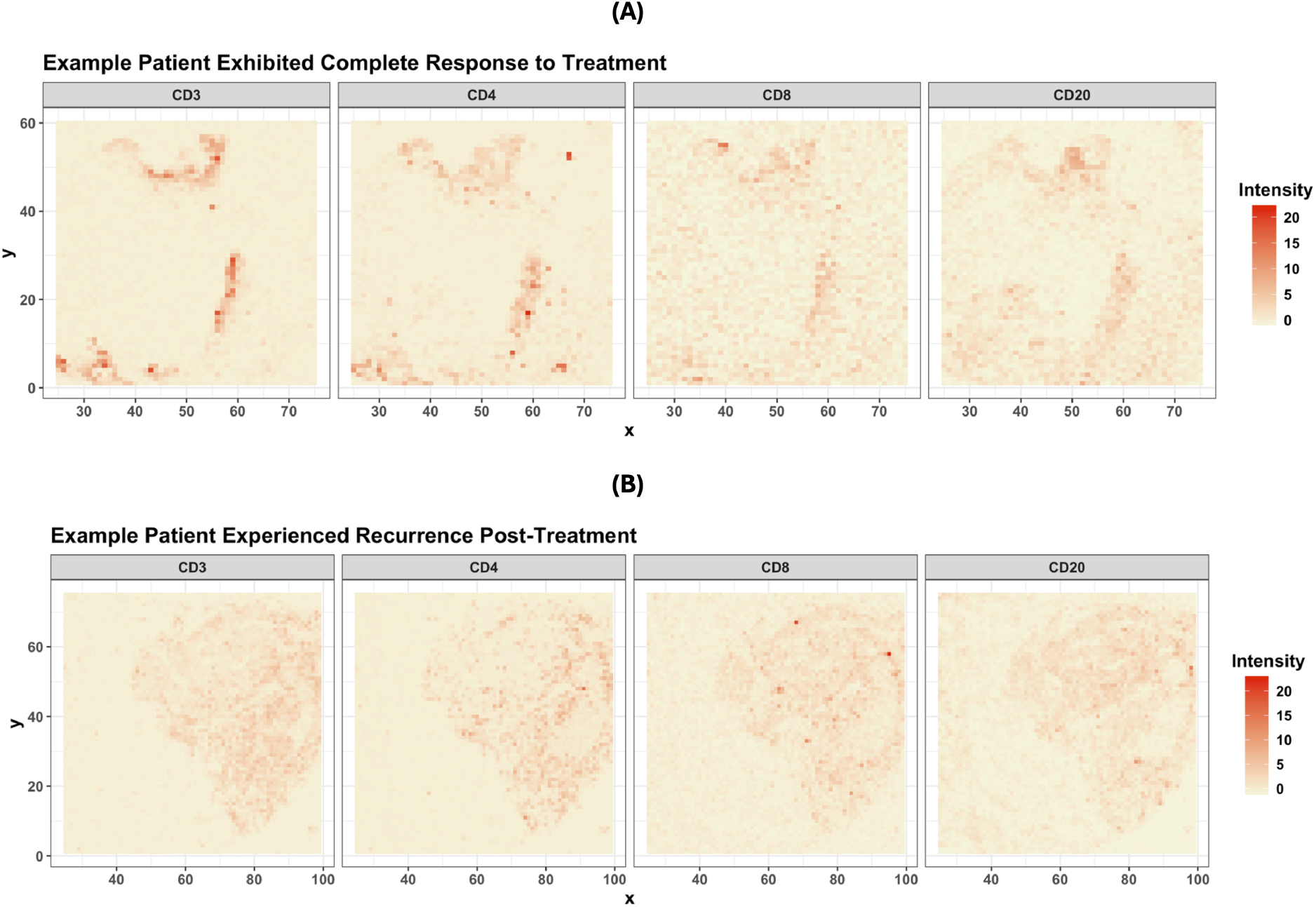
Multichannel TIFF images from two patients from a study of combination chemotherapy plus immunotherapy treatment of triple negative breast cancer. (A) shows the smoothed expression of CD3, CD4, and CD8 protein in a patient who exhibited pathologic complete response (pCR) to combination chemotherapy and immunotherapy treatment. (B) shows the smoothed expression of these proteins in a patient who exhibited recurrent disease (RD) after combination therapy.

We analyzed 238 images across 106 patients. To discretize the topological event history curves to the same grid, we calculated 100 quantiles between 0.1 and 0.9 of the expression of CD3, CD3+CD4, CD3+CD8, and CD20 marker expression across images. For some patients, there were multiple images taken across different regions of the same tumor. To include all images in our analysis, we averaged the estimated curves across images obtained from the same patient to yield a single curve. We then associated these averaged curves with treatment response. For the association test, we considered three approaches: functional modeling, gridwise testing, and kernel testing.

The resulting topological event history curves are shown in Figure 4. Across all four protein combinations, we found that the spatial protein expression within the tumors of patients who experienced pCR on combination chemotherapy plus immunotherapy exhibited fewer births of connected components during filtration. This suggests that protein expression was more concentrated into clusters than in patients who experienced disease recurrence. For CD3+CD4 and CD3+CD20, we found the average curves between the two treatment response groups were most distinct across all quantiles of shared protein expression. These two protein combinations also showed the strongest associations with treatment response across outcome models (Table 4).

**Table 4.** P-values for the association between topological event history curves and response to combination treatment under different outcome models (functional models, gridwise testing, and kernel association testing.)

| Proteins | Functional | Gridwise | Kernel |
| --- | --- | --- | --- |
| CD3 | 0.069 | 0.007 | 0.042 |
| CD3+CD4 | 0.004 | 0.003 | 0.004 |
| CD3+CD8 | 0.075 | 0.030 | 0.146 |
| CD3+CD20 | 0.047 | 0.008 | 0.047 |

**Figure 4.**
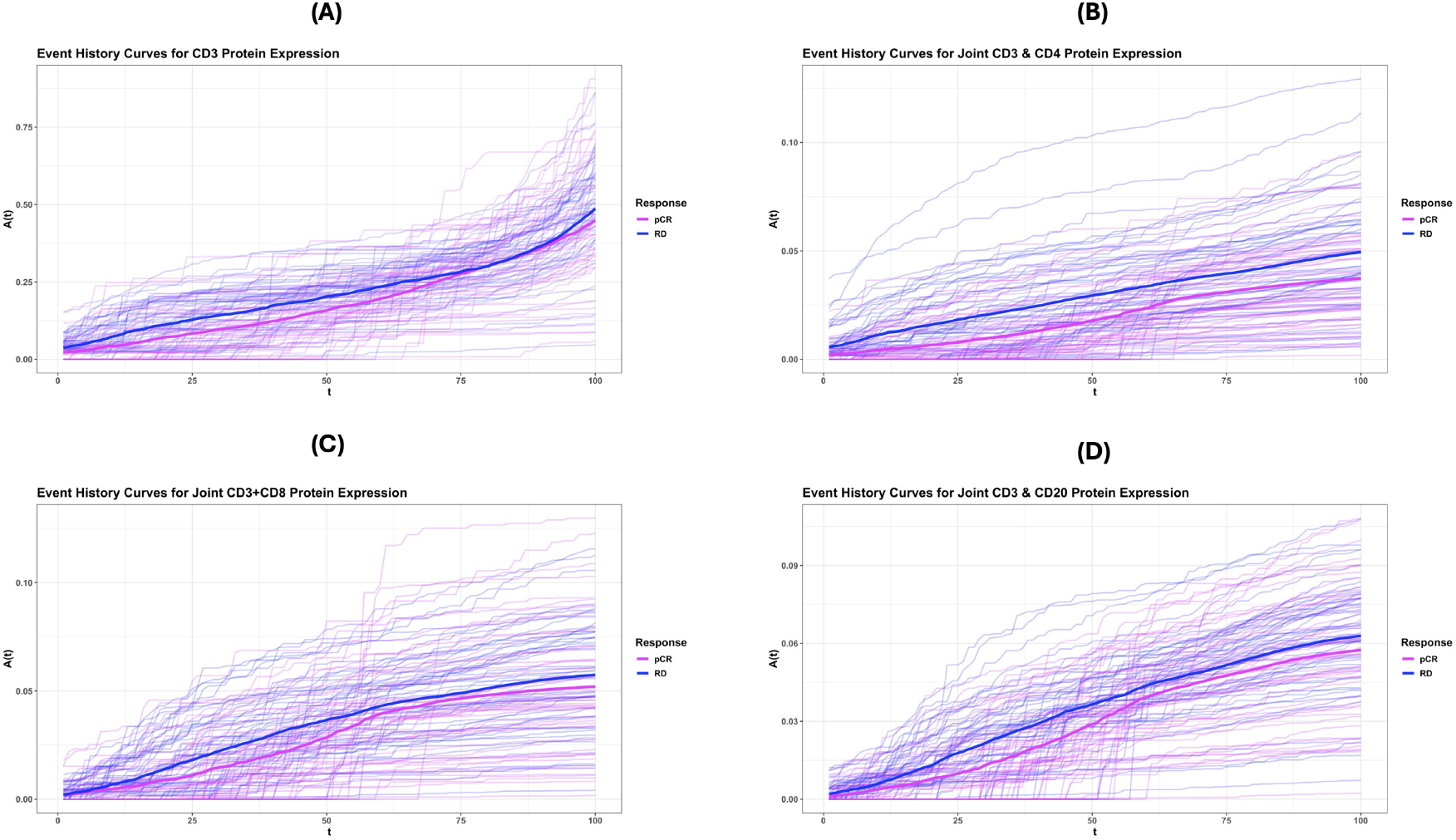
Topological event history curves for the birth of connected components in spatial CD3 expression (A), joint CD3+CD4 expression (B), joint CD3+CD8 expression (C), and joint CD3+CD20 expression (D). Each faded curve reflects the topological event history curve for a single patient. The color of the curve reflects whether the patient experienced pathologic complete response (pCR) or recurrent disease (RD) on combination therapy. The bold curves reflect the average across all patients within each treatment response group (pCR vs. RD).

For some protein combinations, the different outcome modeling approaches offered the same interpretation of the results at a significance level of *α* = 0.05. However, this was not the case for the CD3 protein or the joint expression of CD3+CD8. For CD3, the gridwise and kernel approaches yielded significant p-values (gridwise: *p* = 0.0007, kernel: *p* = 0.042), whereas the functional approach yielded a p-value of 0.069. For CD3+CD8, the gridwise approach yielded a significant p-value (*p* = 0.030) whereas the FPCA and kernel approaches did not (functional: *p* = 0.075, kernel: *p* = 0.146). For these protein combinations, the average topological event history curves either crossed or were closer across quantiles in comparison to CD3+CD4 and CD3+CD20. The gridwise result may have been more significant because the Cauchy combination test was influenced by a small number of very low p-values where the average curves were most distinct. The functional approach, on the other hand, included all functional principal components in the outcome model, some of which may not have described differences between patients who experienced pCR vs. RD. As a result, this may have skewed the test to be insignificant in both cases.

## 5 Discussion

Imaging-based multiplexed spatial proteomics is used to capture the spatial proteome within tissue samples, which may correlate with patient-level outcomes. Existing methods for testing the association between spatial protein expression and outcomes rely on a complex pre-processing pipeline to identify and phenotype individual cells in each image. Segmentation and phenotyping results may be uncertain, and this uncertainty is unaccounted for in downstream analyses. These pre-processing steps also lead to information loss because protein expression beyond the predicted boundaries of cells is ignored in subsequent analyses. We propose a different approach to testing this association which relies on the spatial protein expression itself, circumventing the need to use segmented and phenotyped cell-level data. Our proposed approach, `TOASTER`, uses topological data analysis to characterize the geometric structure of spatial protein expression. The topological structure of each image is summarized using a Nelson-Aalen cumulative hazard function which can then be incorporated into association test using either a functional modeling, gridwise testing, or kernel association testing approach, all of which we explored here.

We found that `TOASTER` yielded a substantial gain in power when analyzing a single protein and a moderate gain in power when analyzing two proteins compared to existing methods. However, it was challenging to properly compare `TOASTER` to existing methods since few have been developed to analyze continuous, spatial protein expression. Nonetheless, we also observed that tracking the births of connected components offered similar power to tracking the deaths of loops and that all three outcome testing approaches yielded comparable power in our simulations. In our analysis of a study in triple negative breast cancer, we found that the topological structure of several protein combinations were associated with treatment response. However, depending on the protein combination, we obtained slightly different results based on the outcome model.

There are several ways to expand this work. First, we addressed the context of analyzing a single protein marker or two protein markers, though one may need to explore higher-order combinations, such as the shared topological structure of CD3, CD4, and CD8 proteins. Our approach may be extended to this context, but, in its current form, as the number of markers increases we expect the number of shared local minima to decrease, which reduces power. To address this, one could consider using a recurrent events framework to characterize the births of topological features during filtration. We also did not consider formally evaluating the best approach for combining topological event history curves across images obtained from the same tissue sample, a common issue in multiplexed imaging [30]. In our analysis of triple negative breast cancer data, we simply took an average across images, though there may be other approaches that improve power, such as by using a weighted average based on a bootstrap variance of each curve [18].

## 6 Data Availability

The data used in our analysis of the NeoTRIP study was obtained via Zenodo at https://zenodo.org/records/7990870.

## 7 Code Availability

An R package to implement `TOASTER` is available at https://github.com/sarahsamorodnitsky/TOASTER.

